# Phospho-signal flow from a pole-localized microdomain spatially patterns transcription factor activity

**DOI:** 10.1101/220293

**Authors:** Keren Lasker, Alex von Diezmann, Daniel G. Ahrens, Thomas H. Mann, W. E. Moerner, Lucy Shapiro

## Abstract

Selective recruitment and concentration of signaling proteins within membrane-less compartments is a ubiquitous mechanism for subcellular organization. However, little is known about how such dynamic recruitment patterns intracellular signaling and cellular development. Here, we combined transcriptional profiling, reaction-diffusion modeling, and single-molecule tracking to study signal exchange in and out of a microdomain at the cell pole of the asymmetrically dividing bacterium *Caulobacter crescentus.* Our study revealed that the microdomain is selectively permeable, and that each protein in the signaling pathway that activates the cell fate transcription factor CtrA is sequestered and uniformly concentrated within the microdomain or its proximal membrane. Restricted rates of entry into and escape from the microdomain enhance phospho-signaling, leading to a sublinear gradient of CtrA~P along the long axis of the cell. The spatial patterning of CtrA~P creates a gradient of transcriptional activation that serves to prime asymmetric development of the two daughter cells.

## INTRODUCTION

The ability of a cell to recruit and coalesce biochemically-related components at specific subcellular sites is crucial for achieving functional complexity and different cell fates across all kingdoms of life (Misteli, 2007; Rudner and Losick, 2010; Shapiro et al., 2009; Wodarz, 2002). While the mechanisms controlling the formation and composition of subcellular domains are increasingly understood, less is known about the spatial and temporal implications of reactions taking place within them. In *Caulobacter crescentus*, the disordered polar organizing protein PopZ establishes a space-filling ~100-200 nm microdomain adjacent to the cell pole that is not encapsulated by a protein shell or a membrane (Bowman et al., 2010; Bowman et al., 2008; Ebersbach et al., 2008; Gahlmann et al., 2013). Here we show that this polar microdomain selectively recruits members of the phospho-signaling pathway that culminates in the activation of the master transcription factor CtrA. We further provide a mechanism for how the concentration and motion of these proteins within the microdomain regulates spatially constrained gene expression and the establishment of asymmetry in the predivisional cell.

*Caulobacter* divides to produce two morphologically distinct daughter cells: a sessile stalked cell that replicates its chromosome following division, and a motile swarmer cell in which chromosome replication is delayed until it differentiates into a stalked cell (Figure 1A, right). This developmental asymmetry is governed by differentially-localized groups of signaling proteins at the two cell poles (Lasker et al., 2016). The localization of these signaling proteins is dependent on the polar organizing protein PopZ that self-assembles into a branched network of filaments *in vitro* and creates a space-filling microdomain at the cell poles *in vivo* (Bowman et al., 2010; Bowman et al., 2008; Gahlmann et al., 2013). PopZ directly binds at least nine proteins that temporally and spatially regulate cell cycle progression, and deletion of the *popZ* gene leads to delocalization of these proteins and to severe cell cycle defects (Bowman et al., 2010; Bowman et al., 2008; Ebersbach et al., 2008; Holmes et al., 2016; Ptacin et al., 2014). Among these binding partners are two signaling proteins that together play a key role in controlling the levels and activity of the master transcription factor CtrA (Quon et al., 1996): the membrane-bound hybrid histidine kinase CckA, which acts as the phosphate source for CtrA, and the small cytoplasmic protein ChpT (Biondi et al., 2006), which shuttles phosphate from CckA to the master transcription factor CtrA (Figure 1A). In its active phosphorylated form (CtrA~P), CtrA controls the transcription of over 100 genes, including those that are necessary for the formation of the nascent swarmer cell, including the flagellar and chemotaxis transcriptional hierarchy (Figure 1A) (Laub et al., 2002; Laub et al., 2000). CtrA~P also serves to inhibit the initiation of DNA replication in the swarmer cell by binding to the chromosome origin (Quon et al., 1998). CtrA levels and activity vary as a function of the cell cycle. In swarmer cells, high CtrA~P levels promote the swarmer fate. During the swarmer to stalk cell transition CtrA~P is cleared from the cell to allow initiation of DNA replication (Domian et al., 1997). In predivisional cells, CtrA proteolysis ceases and its synthesis and activation resumes. Upon cell compartmentalization but prior to division, CtrA~P is proteolyzed in the stalked compartment, while active CtrA~P remains in the swarmer compartment.

**Figure 1.**
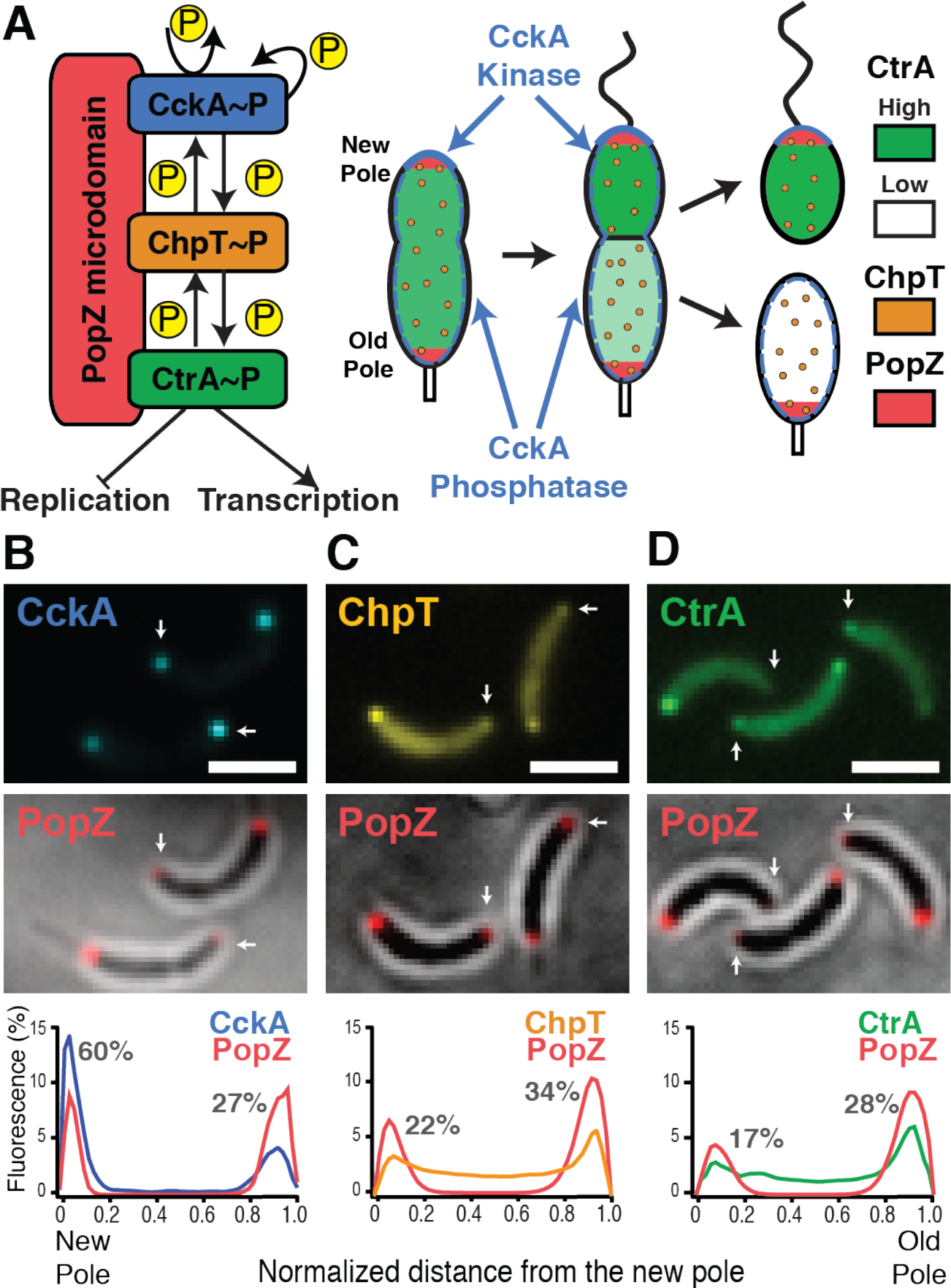
A phospho-signaling cascade sequestered within the PopZ microdomain at the new cell pole culminates in the activation of the cell fate transcription factor CtrA. **(A)** The spatially sequestered phospho-signaling cascade. (Left) The hybrid histidine kinase CckA (blue) autophosphorylates upon localization at the new cell pole (Iniesta et al., 2010; Jacobs et al., 2003; Mann et al., 2016; Tsokos et al., 2011) and subsequently transfers phosphate through the phosphotransfer protein ChpT (orange) to the master transcription factor CtrA (green) (Biondi et al., 2006). Phosphorylated CtrA (CtrA~P) directly controls the transcription of more than 100 cell cycle regulated genes and inhibits the initiation of DNA replication (Laub et al., 2000; Quon et al., 1996; Zhou et al., 2015). Both CckA and ChpT weakly bind to the disordered PopZ protein (Holmes et al., 2016) that self assembles in a space-filling polar hub (Gahlmann et al., 2013) (red). (Right) In the predivisional cell, CckA is positioned at both the new and old cell poles. CckA acts as a kinase exclusively at the new pole to activate CtrA and as a phosphatase away from the new pole (Mann et al., 2016). Upon compartmentalization prior to division, CtrA is degraded in the stalked compartment and remains phosphorylated (and active) in the incipient swarmer compartment (Domian et al., 1997), yielding distinct transcriptional profiles for the two daughter cells. **(B-D)** Diffraction limited images of *Caulobacter* expressing CckA-eYFP (B), ChpT-eYFP (C), or CtrA-eYFP-14 (D) (top row), with accompanying PAmCherry-PopZ images (middle row), new poles marked with white arrow. The fluorescent intensities of PAmCherry-PopZ and each eYFP-labeled protein are plotted versus normalized cell length for a typical cell (bottom row). The percentages of fluorescent signal at the new and old poles are indicated in dark gray for the three CtrA pathway proteins. CckA, ChpT, and CtrA fluorescence intensity profiles were averaged based on 55, 129, and 143 predivisional cells, respectively. Scale bar for B-D is 2 μm.

CckA’s auto-kinase activity is density-dependent and occurs only when CckA is concentrated at the new pole microdomain of the predivisional cell, where it drives phosphorylation of CtrA through ChpT (Iniesta et al., 2010; Jacobs et al., 2003; Jacobs et al., 1999; Mann et al., 2016; Tsokos et al., 2011). In contrast, CckA acts as a phosphatase everywhere else in the cell with density-independent activity (Mann et al., 2016). The antagonism between CckA acting as a kinase and as a phosphatase resembles the process by which concentration gradients are formed in much larger eukaryotic cells and tissues (Wartlick et al., 2009).

For the cytosolic ChpT phosphotransfer protein to interact with CckA at the membrane of the new pole, it must pass through the space-filling PopZ microdomain. ChpT~P must then phosphorylate CtrA to complete the signaling pathway. Thus, critical questions are if (and how) the PopZ microdomain influences access of ChpT to the CckA kinase, where ChpT~P encounters CtrA, and ultimately, what spatial distribution of CtrA~P is established as it emanates from the new pole microdomain. With spatially resolved transcriptional measurements as a proxy for CtrA~P activity, we demonstrate that predivisional cells exhibit a sublinear (approximately exponential) gradient of activated CtrA~P emanating from the new cell pole. By imaging the single-molecule trajectories of CckA, ChpT and CtrA relative to the super-resolved PopZ microdomain, we show that in addition to acting as a localization factor, the microdomain sequesters each member of the CtrA phosphotransfer pathway and restricts the rate at which molecules within the microdomain exchange with the rest of the cell, while restricting polar access for other molecules. Integrating this knowledge with other biochemical data in a comprehensive reaction-diffusion model, we show that both high concentration and slow turnover at the poles are necessary to generate a gradient of CtrA~P activity, in quantitative agreement with our transcriptional measurements. These results demonstrate that a bacterial microdomain adjacent to the cell pole selectively sequesters a phospho-signaling pathway to establish a spatial trajectory of information transfer that controls chromosome readout.

## RESULTS

### The CtrA activation pathway is sequestered within the PopZ microdomain

To understand the influence of the PopZ microdomain on the CtrA activation pathway, we began by determining to what extent the pathway’s proteins were colocalized with PopZ at the old and new poles of predivisional cells. We replaced native *popZ* with an allele for the (photo-activatable) fluorescent protein fusion *PAmCherry-popZ* (Figure S1A, Tables S1-S3). We observed that PopZ forms foci at both poles in predivisional cells as previously shown (Bowman et al., 2008; Ebersbach et al., 2008), with 55% of its signal localized to the old pole and 31% to the new pole on average (Figures 1B-D and S1B). Using strains expressing enhanced yellow fluorescent protein (eYFP) fused to either CckA, ChpT, or CtrA, we measured the co-localization of each pathway component with respect to PAmCherry-PopZ (Figures 1B-D at diffraction-limited 200 nm resolution, also see Figures 3 and 4). Consistent with previous studies (Ebersbach et al., 2008), we found that CckA-eYFP co-localized with PopZ at both the new pole and the old pole, with 60% of total CckA fluorescence intensity localized to the new pole (Figure 1B). While CckA clearly associates with PopZ (Holmes et al., 2016), CckA exhibits a greater concentration at the new pole due to additional pole-specific localization factors (Iniesta et al., 2010). The cytosolic phosphotransferase ChpT, like its interacting membrane histidine kinase CckA, was shown to bind PopZ *in vivo* using the heterologous *Escherichia coli* system (Holmes et al., 2016). We thus speculated that PopZ-binding interactions would also cause ChpT to be enriched within the PopZ microdomain in *Caulobacter*. Indeed, ChpT formed foci at both poles of the cells (Figures 1C and S1B). While ChpT weakly binds CckA *in vitro* (Blair et al., 2013), ChpT did not show greater localization at the new pole, where most of CckA molecules reside. These results suggest that PopZ is a critical polar localization factor for ChpT in *Caulobacter*.

Unlike ChpT and CckA, CtrA does not colocalize with PopZ when heterologously expressed in *E. coli*, suggesting that binding interactions between CtrA and PopZ are weak or nonexistent (Holmes et al., 2016). We asked whether CtrA forms a focus within the PopZ microdomain in *Caulobacter* without direct binding to PopZ, as does the cytosolic protein DivK (Holmes et al., 2016; Lam et al., 2003). Previous imaging of an N-terminal YFP fusion to CtrA in predivisional cells showed a large diffuse population of CtrA with transient accumulation at the old cell pole prior to proteolysis, independent of CtrA’s phosphorylation state or its degradation motif (Ryan et al., 2004), while no CtrA accumulation was detected at the new pole. Because the N-terminal domain of CtrA is important for function, we repeated this experiment using a sandwich fusion that integrated eYFP between CtrA’s DNA binding domain and its 14-residue C-terminal degradation tag (Figure S1A) (CtrA-eYFP-14). We observed a localization profile of CtrA-eYFP-14 in synchronized predivisional cells (Figure S1C) in which 17% of the fluorescent signal was at the new pole and 28% of the signal was located at the old pole. Altogether, our imaging results show that all three members of the phospho-signaling pathway are greatly enriched within the PopZ microdomains of the *Caulobacter* cell.

### CtrA exhibits a gradient of activity in predivisional cells

We designed a transcription assay as a proxy for measuring the CtrA~P spatial profile and used it to determine whether polar accumulation of the proteins in the CtrA activation pathway regulates the distribution of CtrA~P across the cell. We introduced a CtrA~P activated promoter (P_*350*_, from the CCNA_00350 operon) driving an *eyfp* gene at four different sites along the right arm of *Caulobacter’*s single circular chromosome (loci L1-L4) (Figure 2A, Table S4). Locus L1 is adjacent to the replication origin, which is positioned near the cell pole, while locus L4 is positioned near the chromosome terminus (Figure 2A). Because the position of any given gene on the *Caulobacter* chromosome reflects its position within the cell (Viollier et al., 2004), we can approximate the spatial position of loci L1-L4 within the cell and their time of replication (SI, Table S4).

**Figure 2.**
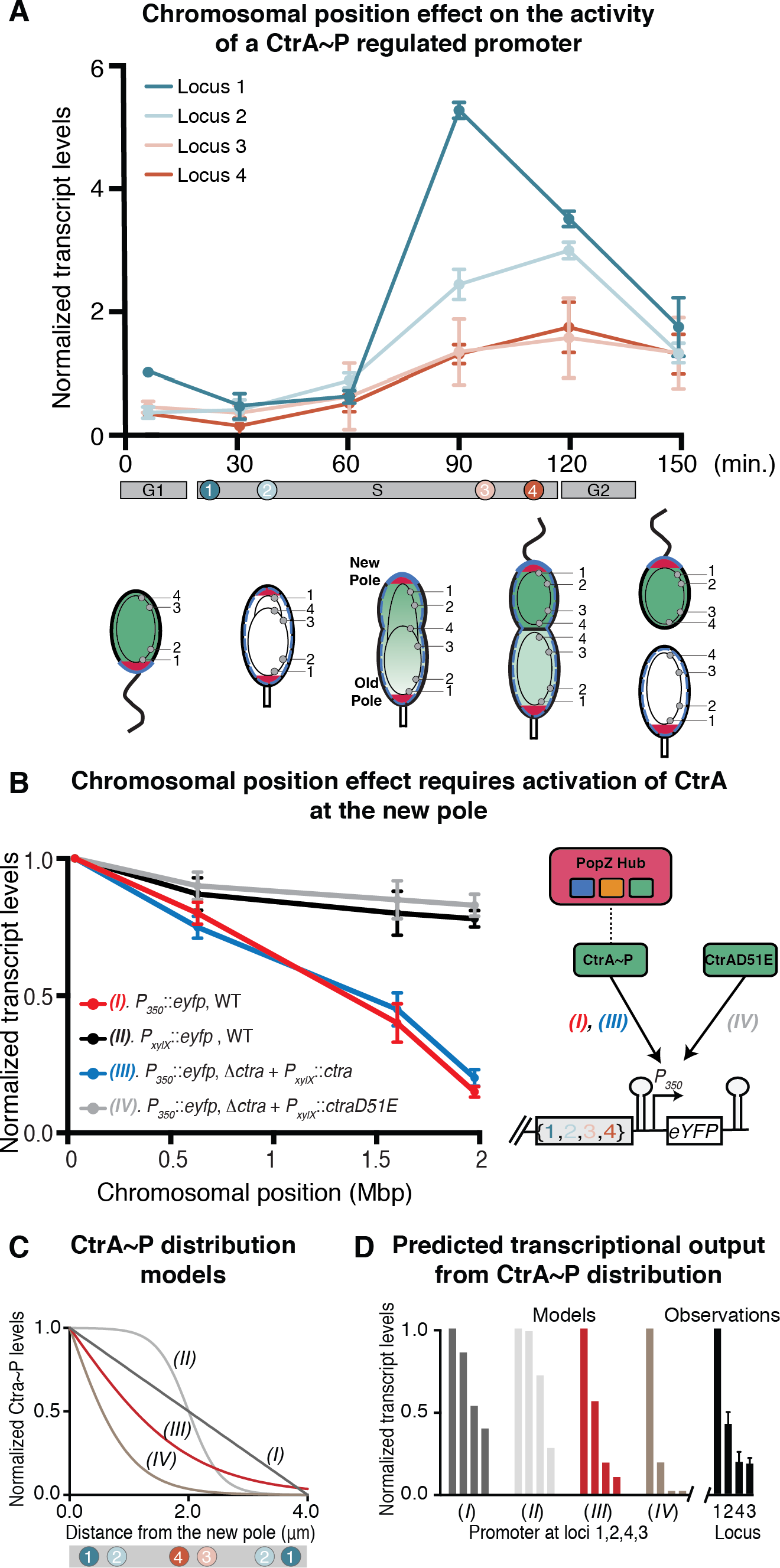
Predivisional cells exhibit a spatial gradient of CtrA~P activity. **(A)** Time resolved RT-qPCR of mRNA from an *eyfp* gene under the control of the CtrA-regulated P_*350*_ promoter integrated at four loci of the chromosome (L1-L4). Transcript levels were assayed every 30 minutes during the 150-minutes cell cycle. A cell cycle schematic (lower) illustrates the four chromosomal loci and the progression of DNA replication and segregation. Replication times of loci L1-L4 are shown below the qPCR time points. The chromosome is represented by black ovals and theta structures within each cell. The red areas of the cell poles indicate the PopZ microdomain and the green tone indicates CtrA~P levels. Transcription is normalized to the five-minutes time point of site 1. Transcription from locus L1 peaked at the 90 minute time point and reached an activity level of 6 AU while transcription from loci L2-L4 peaked at the 120 minutes time point, reaching values of 3, 1.9, and 1.8 AU respectively. **(B)** RT-qPCR of mRNA from an *eyfp* gene integrated at four loci of the chromosome measured in mixed population cells in four different strain backgrounds. Transcription of *eyfp* was driven by either the xylose-inducible *P_xyl_* (black), or by the CtrA~P activated promoter, P_*350*_. WT CtrA was produced from its native locus (red), or on a plasmid in cells lacking chromosomal *ctrA* (blue). The phospho-mimetic *ctrA*(*D51E*) was produced from a plasmid in a background also lacking native *ctrA* (gray). On the right, a diagram shows the signal pathway for the CtrA~P-controlled promoters and for the phospho-mimetic CtrA(D51E)-controlled promoters that are independent of the signaling cascade. Error bars for A and B represent the standard deviation of three independent experiments. **(C)** Four possible profiles of CtrA~P distribution along the long axis of the predivisional cell. Note that CckA acts as a kinase specifically at the new pole microdomain. The positions of loci L1-L4 in the predivisional cell are shown in colors along a gray bar. **(D)** Calculated transcript levels of *eyfp* gene driven by the P_*350*_ CtrA-regulated promoter integrated at chromosomal loci L1-L4 for each of the models (I-IV). Despite the cooperative activity of CtrA~P at P_350_, neither linear (i, dark gray) nor sigmoidal (ii, light gray) CtrA~P profiles recapitulated the measured mRNA production. By contrast, an appropriately steep sublinear distribution (iii, red) best recapitulated the transcriptional output obtained experimentally for RT-qPCR of predivisional cells (black, Figure 2A).

We used reverse transcription quantitative PCR (RT-qPCR) (Heid et al., 1996) to measure the reporter *eyfp* mRNA transcribed from each locus L1-L4 in synchronized populations beginning from swarmer cells (Figure 2A). This allowed us to quantitatively and directly compare the activity of CtrA~P regulated promoters from each spatial position over the course of the cell cycle. For clarity, we report the transcriptional output relative to the 5-minutes time point of L1 (1 AU). Over the course of the experiment, the single chromosome replicates once, doubling the output of each locus as it is replicated. While transcription from all four integration loci remained low during the first 60 minutes of the cell cycle, the loci began to diverge at the onset of CtrA activation in predivisional cells. Critically, at the 90-minutes time point, transcriptional activity at site L1 peaked to 5.5±0.11 AU, while transcription from site 2 peaked to 2.5±0.25 AU, even though both sites had been duplicated far earlier (Figure 2A, Table S4). Transcription from sites L2-L4 peaked at the 120-minute time point, upon completion of DNA replication, with 1.6±0.67 AU and 1.8±0.42 AU for sites L3 and L4 (Figure 2A). Similar patterns of locus-dependent promoter activity in synchronized predivisional cells were obtained for a different CtrA-regulated promoter (*PpilA*, Figure S1E). In contrast, the CtrA-independent xylose-inducible promoter showed only ≤ 2-fold increases consistent with the timing of chromosome duplication (Figure S1D). Further, we note that despite the roughly uniform distribution of CtrA binding motifs across the chromosome, ChIP-seq data show that CtrA molecules bind more frequently overall to the origin-proximal regions of the chromosome (Figure S1F). These results indicate that the activated CtrA~P transcription factor exhibits spatial control of promoter activity, with highest transcription from loci positioned near the newly replicated origin of replication at the new cell pole.

To confirm that CtrA activity was spatially regulated by its phosphorylation state, we decoupled CtrA from the phosphotransfer pathway by deleting the native *ctrA* gene and replacing it with a plasmid-borne, xylose-inducible phosphomimetic mutant, *ctrA*(*D51E*) (Figure 2B). This mutant cannot accept a phosphate, but the glutamate replacing the aspartate results in constitutive CtrA activity, (Domian et al., 1997), decoupling it from the CckA-ChpT phospho-transfer pathway. When *ctrA*(*D51E*) was expressed as the sole copy of CtrA, position-specific effects on transcription from the CtrA-activated P_*350*_ promoter were lost, demonstrating that the gradient in CtrA activity depended on the signaling cascade emanating from the cell pole. This effect was not due to changing protein levels, as xylose-induced expression of a plasmid-borne wildtype *ctrA* still resulted in a gradient of P_*350*_ activity (Figure 2B). Collectively, these experiments indicate that localized activity of the CckA phospho-signaling pathway leads to a gradient in CtrA~P and spatial control of CtrA-controlled promoters.

Next, we used mathematical modeling to relate the measured transcriptional response from the *P_350_* loci to local CtrA~P concentration. CtrA~P dimers bind cooperatively to many promoters (Reisenauer et al., 1999; Siam and Marczynski, 2000). Because CtrA~P binds P_*350*_ as a dimer as well (Zhou et al., 2015), we modeled the transcriptional response from P_*350*_ as a Hill function, with parameters fitted based on previously measured CtrA copy number and cell cycle dependent transcription of *350* (Schrader et al., 2016) (Equation S7, Figure S1G). For comparison, we present the effect of several different example CtrA~P distributions on the transcriptional output from the four integration loci (L1-L4) (Figures 2C-D). Even including the effects of cooperative activity, only a sublinear drop in CtrA~P levels away from the new pole can recapitulate the 2-fold decrease in mRNA levels from L1 to L2 and the 5-fold decrease in mRNA levels from L1 to L3 and L4 (Figures 2C and 2D *III*). Thus, our measurements of transcriptional output at distinct chromosomal loci not only demonstrate the existence of spatially patterned CtrA~P activity, but show that this pattern is sharply peaked at the new pole, proximal to the CckA kinase, and falls dramatically away from the pole.

### CckA molecules are concentrated and their diffusion is slowed within the polar PopZ microdomain

The colocalization of the CtrA pathway proteins at the new pole microdomain (Figure 1) and the formation of a non-linear CtrA~P gradient (Figure 2) suggested a possible means to spatially control CtrA~P activation state: joint recruitment and interaction within this microdomain. To resolve protein localization and dynamics at the poles, we employed single-molecule tracking of the CtrA pathway proteins combined with super-resolution microscopy of the static PopZ microdomain, beginning with CckA. To track CckA motion over the curved inner membrane, we used an engineered point spread function (the DH-PSF) (Gahlmann et al., 2013) to localize and follow the motion of CckA-eYFP molecules in three dimensions (3D), avoiding the 30% errors that would be present for 2D trajectories of membrane proteins (Figure S2A-C).

By correlating our tracking measurements with 3D super-resolution imaging of a xylose-induced copy of PAmCherry-PopZ in a merodiploid background, we precisely defined what parts of the CckA-eYFP trajectories explored the membrane area of the PopZ microdomains at the new and old poles (Figure 3A). Using mean-squared-displacement (MSD) analysis, we found that CckA diffused most rapidly in the cell body, 2-fold slower in the old pole, and 4-fold slower in the new pole (D = 0.0082 ±0.0020, 0.0040 ± 0.0014, and 0.0022 ± 0.0013 μm^2^/s, respectively) (Figure 3B). One certain cause for the reduction in diffusivity at the poles is polar binding interactions, both to PopZ and to CckA-specific localization factors at the new pole (Holmes et al., 2016; Iniesta et al., 2010). Another likely cause is the formation of higher-order CckA assemblies (Mann et al., 2016), which would presumably be enhanced at the high CckA concentrations within the new pole (and to a lesser extent, within the old pole).

**Figure 3.**
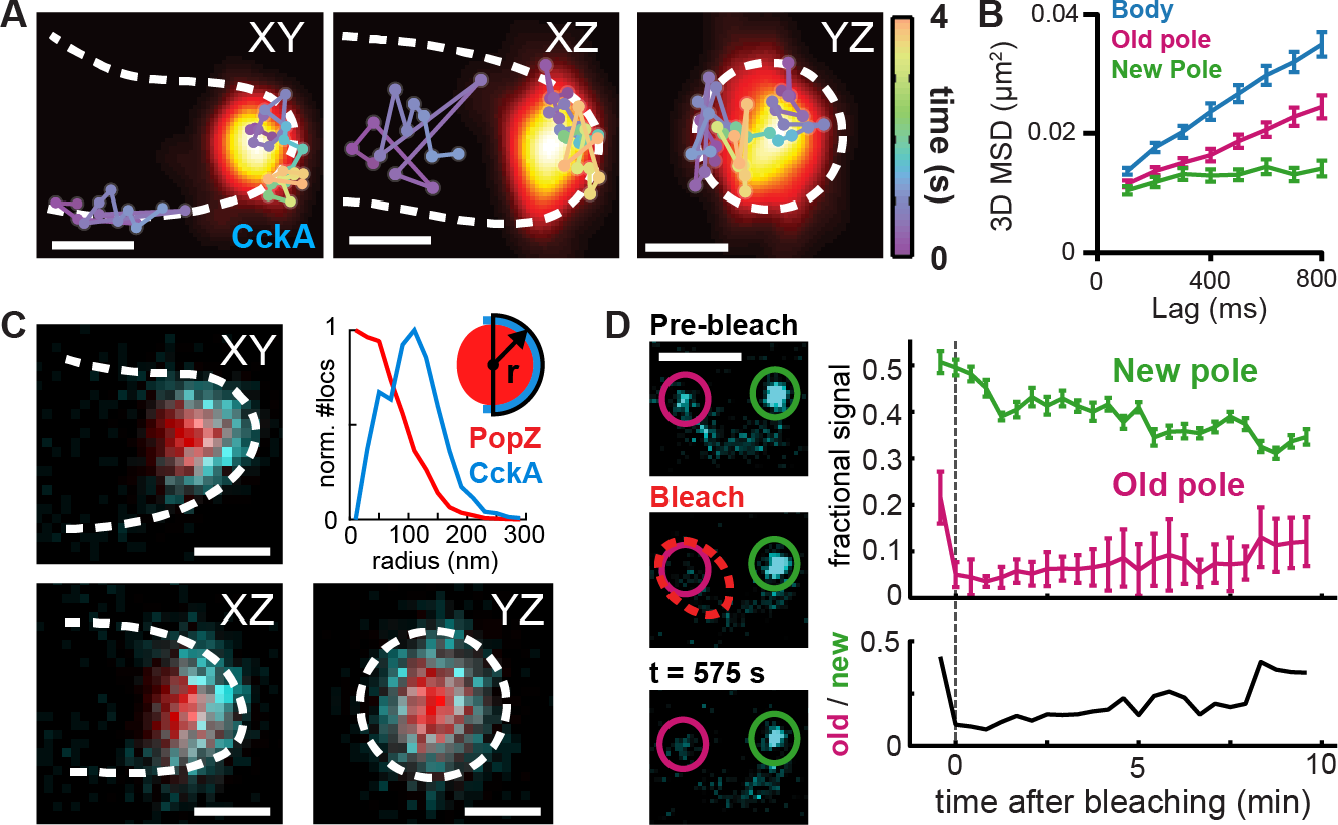
CckA dynamics and distribution within the poles. **(A)** 3D single-molecule tracking of CckA (time-coded connected dots) relative to a superresolution reconstruction of PopZ (yellow-orange). Two example tracks are shown in three different orientations. **(B)** Mean-squared-displacement curves for CckA tracks within selected cellular regions. Errorbars: 95% CI from bootstrap analysis of tracks. **(C)** Average CckA and PopZ polar distributions using 3D localization data from N = 29 old poles (2006 and 5282 localizations respectively). To emphasize the radial distribution of CckA on the membrane, each view shows a 200 nm thick slice (e.g. the XY slice has 200 nm Z thickness). Plot shows the radial distribution of CckA and PopZ from the PopZ centroid with volume-normalized density. **(D)** Loss/recovery from the new and old poles following targeted photobleaching of old pole CckA-eYFP. Signal is defined as the fraction of total cell fluorescence (upper plot, errorbars show ± 95% CI). After 10 minutes, while the total amount of fluorescent CckA-eYFP has been reduced by targeted photobleaching, the initial ratio of old to new pole signal is restored (lower plot). Average of N = 6 cells. Scale bar in **(D)**, 2 μm; all other scale bars, 200 nm.

To define the average nanoscale distribution of CckA at the new and old poles, we combined high-resolution CckA and PopZ localization data from many individual microdomains (SI). We found that CckA molecules were distributed roughly evenly throughout the 3D membrane region adjacent to PopZ, and that the CckA concentration dropped off sharply away from the microdomain (Figure 3C, Figure S2D). While the uniform density in the averaged CckA distribution does not preclude the possibility of individual CckA clusters within microdomains (as has been suggested in PopZ overexpression experiments (Ebersbach et al., 2008)) it clearly indicates that CckA does not have a preference for particular positions within the PopZ microdomain. CckA formed a hemispherical cup surrounding PopZ in old poles (Figure 3C), while in new poles, the PopZ microdomain took up a smaller volume, and CckA occupied a proportionally smaller area on the new pole membrane (Figure S2D). Notably, the area taken up by CckA at the new pole was lower despite having a higher total number of CckA molecules (Figure 1C). From the nanoscale geometry of CckA and calculations of copy number (SI), we estimated a CckA concentration of ~5,000-10,000 and 1,250 molecules/μm^2^ for the new and old poles, respectively, more than 100x greater than the concentration in the cell body (9 molecules/μm^2^). This estimated concentration of CckA at the new pole is close to the previously determined CckA concentration on liposomes *in vitro* that leads to maximum autokinase activity, while the estimated CckA concentration at the old pole was shown *in vitro* to lead to little or no autokinase activity (Mann et al., 2016).

To determine whether CckA was free to enter and exit the pole over a longer timescale than seconds, we used a diffraction-limited confocal microscope to photobleach CckA at the old pole and measured the timescale of fluorescence recovery and loss at the old and new poles (Figure 3D and Figure S3). While targeted photobleaching depleted ~30% of the total cell fluorescence, we found that the ratio of CckA within the new and old poles was restored close to its pre-bleach value after 10 minutes. This was consistent with reaction-diffusion simulations using the experimentally measured diffusivities and assuming transient binding of CckA to species within the microdomain (Figure S3E). Thus, while CckA recovery was slower than would be expected for free diffusion, CckA was still mobile and not irreversibly bound within the polar microdomain. Collectively, the CckA tracking and photobleaching experiments indicate that CckA is concentrated throughout the polar membrane cap of the PopZ microdomain by crowding and reversible binding to other proteins within the microdomain.

### ChpT and CtrA molecules are transiently sequestered within the PopZ microdomain volume

We then turned to the smaller, cytoplasmic members of the phosphotransfer pathway, ChpT and CtrA, to determine to what extent the polar microdomains affected localization and transport of these molecules in predivisional cells. ChpT-eYFP molecules outside the poles diffused rapidly throughout the cell, but were captured within the PopZ volume upon reaching the poles, as clearly shown by trajectories with long polar residence time (Figure 4A). While most captured ChpT molecules diffused throughout the microdomain volume, a fraction of ChpT-eYFP trajectories exhibited motion only in the plane of the polar membrane (Figure 4B), suggestive of binding to CckA. Similar to ChpT, we observed that CtrA-eYFP-14 entered and was slowed within the PopZ microdomain (Figure 4C). CtrA and ChpT molecules exited the microdomain after a short time period (Figure 4D; ChpT not shown) and immediately regained their previous diffusivity within the cell body. Pooling localizations from many cells, we observed that both proteins were recruited roughly uniformly throughout the PopZ volume (Figure 4E and Figure S4A) with a concentration relative to the body comparable to that observed in our diffraction-limited images (Figure 1D,E). As CtrA does not have affinity for PopZ but does bind ChpT (Blair et al., 2013; Holmes et al., 2016), this suggests that ChpT molecules embedded in the microdomain act as binding partners to indirectly recruit CtrA. We also observed a slight enrichment of CtrA at the cytoplasmic face of the microdomain at both the new and old poles, ~50-100 nm from the PopZ centroid (shoulders on green curves in Figure 4E), potentially reflecting binding to the chromosome.

**Figure 4.**
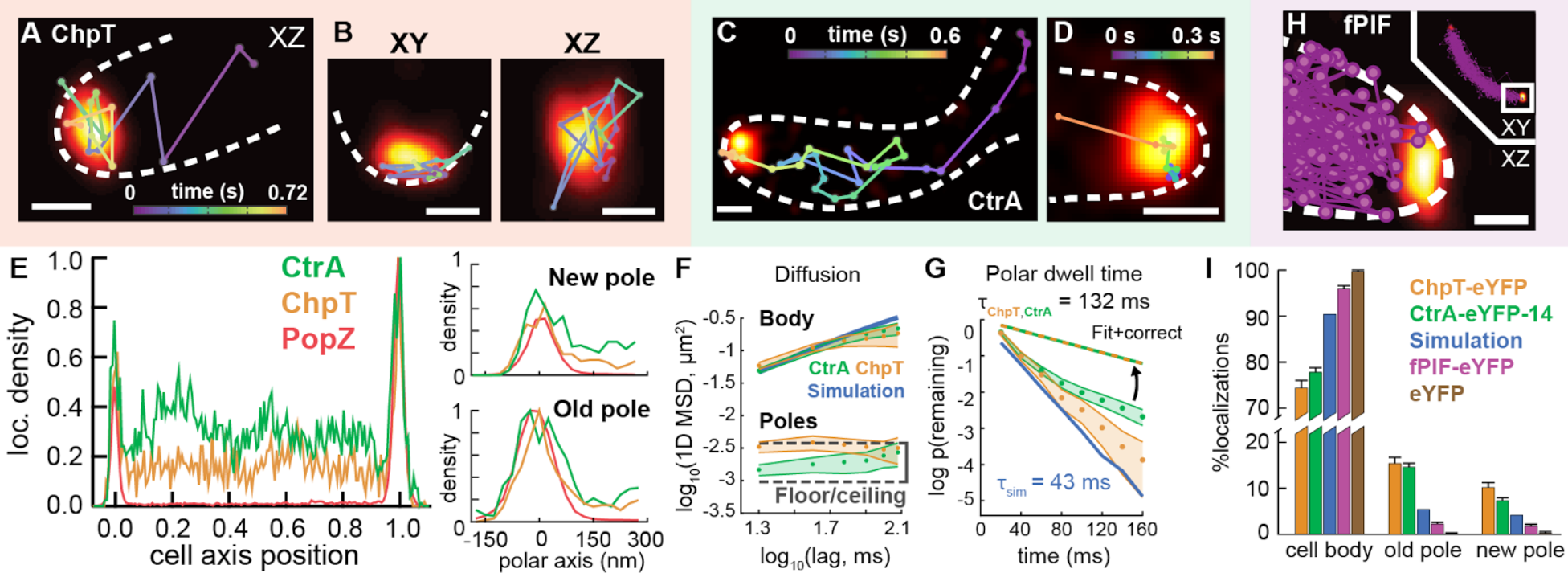
The polar microdomain concentrates and slows ChpT and CtrA, while excluding other small cytoplasmic proteins. **(A-B)** Two cells showing exemplary single-molecule trajectories of ChpT-eYFP (time-coded connected dots) with PAmCherry-PopZ super-resolution reconstructions (yellow-orange). **(A)** ChpT entry into the pole and capture by the PopZ microdomain. This example shows the old pole. **(B)** Two views of the same ChpT molecule as it exhibits apparent membrane-associated motion within the PopZ microdomain at the new pole. **(C-D)** Two cells showing 2D single-molecule trajectories of CtrA-eYFP-14 relative to PAmCherry-PopZ. **(C)** CtrA entering the PopZ microdomain at the new pole. **(D)** CtrA transiting through and leaving the old pole microdomain. **(E)** Average distributions of CtrA (N = 60 cells), ChpT (N = 27), and PopZ (N = 27), along the cell axis and in the local coordinate system of the cell poles. **(F)** Log-log MSD plots of CtrA (green) and ChpT (orange) motion projected along the *Caulobacter* cell axis in both the body and poles. Shaded area: 95% CI from bootstrap analysis of tracks. Blue line: MSD for simulated free diffusion with D = 1.8 μm^2^/s (Figure S6). Floor/ceiling: the minimum and maximum MSD values within the pole, corresponding to static molecules and molecules that explore the pole within a frame. **(G)** The decay rates of ChpT and CtrA escaping from the pole or bleaching before escape. Shaded area: 95% CI from bootstrap analysis of tracks. Linearity of the cumulative distribution function of survival, p(remaining), on the semi-log plot indicates an exponential distribution (Distributions from N = 434 and 1149 events of which 77.1% and 80.9% were due to bleaching, respectively.) Blue line: distribution of escape rates from the pole for simulated molecules diffusing with D = 1.8 μm^2^/s without hindrance by PopZ (Figure S6). Exponential fit lifetimes (1/e) shown. **(H)** Representative cell displaying an overlay of 92 total 3D single-molecule tracks of fPIF-eYFP shown relative to a PAmCherry-PopZ super-resolution reconstruction. **(I)** The proportion of localizations within the cell body, the new pole, and the old pole, as observed for tracks of ChpT-eYFP, CtrA-eYFP-14, fPIF-eYFP, and eYFP, and for simulated diffusing molecules unconstrained by PopZ (Figure S6). Magnitude of 95% CI shown as lines above data. Scale bars: 200 nm.

We used MSD analysis to quantify the diffusivity of ChpT and CtrA, projecting motion onto the (curved) 1D cell axis to avoid confinement effects arising from the narrow cell width (Figure 4F). Both ChpT and CtrA exhibited anomalous subdiffusive motion in the cell body, apparent as a slope less than 1 in the log-log MSD plot. Given the short length of our eYFP trajectories, fits to these curves only allowed us to give an upper bound for the subdiffusive exponent α, indicating α < 0.7 and a short-timescale apparent diffusivity *D_app_* = 1.8 μm^2^/s (SI, Figure S4H). We also observed anomalous subdiffusion in trajectories of free eYFP, which we do not expect to directly bind any targets in *Caulobacter* (Figure S4). Thus, subdiffusion in the cell body likely results from general properties such as crowding in the cytoplasm and obstruction by the nucleoid.

Within the poles, both ChpT and CtrA exhibited MSD values an order of magnitude lower than in the cell body (Figure 4F). The MSD of ChpT molecules within the pole reached an asymptotic limit of ~10^−2.5^ μm^2^ (3x greater than our 24 nm localization precision) independent of lag. This result was consistent with close to complete exploration of the 150-200 nm polar volume within a 20 ms frame (SI), implying D ≥ 0.1 μm^2^/s. In contrast, polar CtrA exhibited a steady increase of its MSD with lag time between the limits of error and free diffusion, as expected from our observation of tracks that slowly explored the PopZ microdomain (Figure 4D), and fits to these data gave D ≈ 0.01 μm^2^/s (Figure S4). Thus, both ChpT and CtrA were relatively free to explore the polar space in which they were concentrated.

While ChpT and CtrA appeared to move freely within the PopZ microdomain, MSD analysis alone cannot determine to what extent molecules are hindered from leaving the pole. To measure this quantity, we calculated the dwell times of eYFP-labeled molecules within the polar microdomain before returning to the cell body (Figure 4G) and found that most molecules exited the microdomain on the ≤ 100 ms scale (i.e. generally faster than the exemplary trajectories shown in Figure 4A-D). The apparent exit rate was increased due to competition between true exit events and the rapid photobleaching of eYFP labels. We compensated for this effect by approximating photobleaching and exit as competing exponential processes, scaling the rates by the proportion of tracks that exited before photobleaching (SI). Using this approximation, we measured similar dwell times of ChpT and CtrA within the poles: 132±39 ms and 132±28 ms (95% CI), respectively, as shown by the corrected survival curve for both. This was three-fold slower than the rate of exit in simulations of free diffusion using *D_app_* = 1.8 μm^2^/s, indicating that slowed polar diffusion due to binding and/or physical obstruction within the crowded PopZ microdomain acts to slow protein turnover at the poles.

### The PopZ microdomain creates a barrier to entry for non-client proteins

Ribosomes and chromosomal DNA do not enter the PopZ microdomain, an effect ascribed to such large molecules being unable to pass through the fine pores of a filamentous PopZ matrix (Bowman et al., 2010; Ebersbach et al., 2008). We wondered whether the slowed diffusion and escape rates of ChpT and CtrA within the PopZ microdomain was due in part to percolation through such a matrix. To isolate this effect, we performed single-molecule tracking of proteins heterologous to *Caulobacter*, for which binding interactions to polar localization factors should be minimal. We selected two proteins: eYFP by itself, and eYFP fused to a 100 amino acid fragment of *A. thaliana* PIF6 which we term fPIF (Levskaya et al., 2009). As eYFP was 0.5 times and fPIF-eYFP was 0.75 times the mass of ChpT-eYFP and CtrA-eYFP-14 (Table S1), as reflected by their faster diffusion in the cell body (Figure S4), we predicted that both proteins would penetrate the PopZ matrix. Yet surprisingly, both eYFP and fPIF-eYFP only explored the 3D volume of the cell up to the edge of PopZ, and did not enter the microdomain (Figures 4H and S4E-F).

We quantified the polar recruitment and exclusion of ChpT, CtrA, fPIF, and eYFP by counting single-molecule localizations that appeared within the super-resolved reconstruction of PopZ, and comparing these patterns to the expected distribution in the absence of microdomain interactions from simulations of free diffusion throughout the cell volume (Figures 4I and S4). Relative to the “free diffusion” case, ChpT and CtrA were concentrated ~3x at the old pole and ~2x at the new pole, reflecting their direct (PopZ-binding) and indirect (ChpT-binding) recruitment. By contrast, fPIF and eYFP were not only not enriched, but were actively excluded from the poles; while a small fraction (4.0 ± 0.6%) of fPIF molecules were scored as polar, these molecules were generally located at the cytoplasmic interface of the microdomain, not the interior (Figure 4H). As fPIF and eYFP were excluded despite being smaller than ChpT and CtrA, pore size cannot explain this effect. In contrast to our experiments with wildtype cells, fluorescent proteins are able to enter the PopZ microdomain when PopZ is heterologously expressed in *E. coli* (i.e., in the absence of known clients), or when PopZ is enlarged 10-20 fold to 2-4 μm in *Caulobacter*, greatly increasing the available volume (Ebersbach et al., 2008; Laloux and Jacobs-Wagner, 2013). This suggests that in addition to the presence or absence of specific binding interactions to PopZ or other polar proteins, general properties such as the volume and client occupancy of the PopZ microdomain may also control the microdomain permeability in a nonspecific way.

### CtrA activation occurs within and is modulated by the new pole microdomain

The dense recruitment and colocalization of the CtrA pathway proteins within the PopZ microdomain suggested a possible mechanism for our finding that CtrA~P sharply decreases away from the new pole in the predivisional *Caulobacter* cell. To quantitatively investigate how concentration and restricted motion within the microdomain patterns CtrA~P amplitude and distribution, we developed a continuum reaction-diffusion model. This model lets us compute the concentrations and phosphorylation states of CckA, ChpT, and CtrA as a function of diffusion and biochemical interactions inside and outside of the PopZ microdomains (see Figures S5A-B, Table S6 and SI for details). Our high-resolution, quantitative measurements of protein localization profiles let us precisely define the binding coefficients of CckA, ChpT, and CtrA at the pole and ensured that our model correctly recapitulates the true molecular concentrations *in vivo* (Figures 1B-D, 4E, and S5C). Using measured data on phosphotransfer rates *in vitro* let us calculate the steady-state phosphorylation level of each protein within new cell pole microdomain (93% of CckA~P, 68% of ChpT~P, and 10% of CtrA~P). The ChpT~P concentration sharply dropped from the new pole microdomain and continued to decline with a shallower sublinear gradient (Figure 5A). In contrast, the CtrA~P concentration declined smoothly away from the new pole microdomain with a sublinear distribution (Figure 5A). Following our calculation of transcriptional response from the *P_350_* promoter integrated at the four chromosomal loci (Figures 2C and 2D), we calculated transcriptional output in response to the modeled CtrA~P distribution. The calculated *eyfp* mRNA values recapitulated the measured mRNA levels in our model (Figure S5D). Thus, our model demonstrates that the dense localization of the phosphotransfer pathway can be expected to give rise to the observed sharp gradient of CtrA~P in predivisional cells.

**Figure 5.**
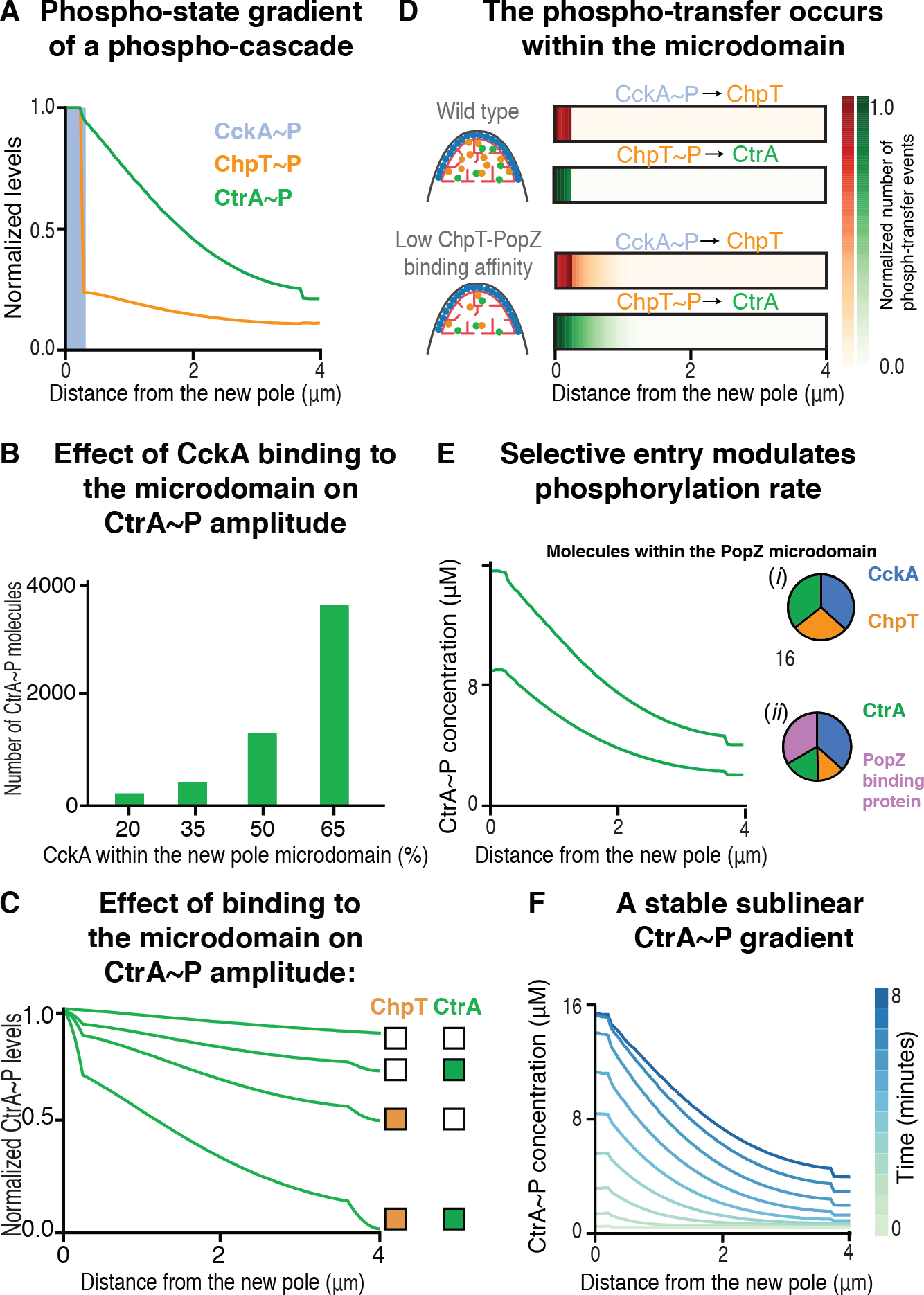
Sequestration within the PopZ microdomain creates an activity gradient of CtrA. **(A)** The predivisional cell maintains a gradient of ChpT~P and CtrA~P. Steady state distributions of CckA~P, ChpT~P, and CtrA~P (light blue, orange, and green) as derived from our reaction-diffusion model incorporating measured diffusion coefficients (Figures 3 and 4). The phosphorylated forms of all three proteins are localized to the new pole microdomain, where CckA acts as a kinase. Both ChpT~P and CtrA~P exhibit a gradient descending from the new pole. CtrA~P forms a steep sublinear gradient, such that most CtrA~P molecules reside in the nascent daughter swarmer cell compartment. Phosphotransfer from ChpT~P to form CtrA~P occurs rapidly within the pole, depleting most ChpT~P molecules before exit and causing the relative levels of ChpT~P in the body to be much lower than those of CtrA~P, although still maintaining a shallow gradient. In comparison to the phospho-gradients of ChpT and CtrA, the total populations of ChpT/ChpT~P and CtrA/CtrA~P within the cell body were evenly distributed (Figure S7A). (B) Steady-state levels of CtrA~P as a function of CckA accumulation at the new pole microdomain. (C) CtrA~P distribution as a function of new pole accumulation of ChpT and CtrA. Steady-state distribution of CtrA~P in four conditions altering the polar accumulation of ChpT and CtrA. In these four conditions ChpT and CtrA are either uniformly distributed (empty boxes) or have 40% total protein accumulated at the two poles (filled boxes). (D) Phosphotransfer events occur at the new pole microdomain. “Phospho-graphs” showing the number of CckA~P to ChpT and ChpT~P to CtrA phosphotransfer events at steady state. The phosphotransfer events occur exclusively within the new pole microdomain in a simulation using our measured parameters (upper phospho-graph). However, in an *in-silico* mutant with weak binding between ChpT and the PopZ microdomain, phosphotransfer events are dispersed across the cell (lower phospho-graph). (E) Introduction of a competitor for PopZ binding sites in the new pole microdomain decreases the polar concentration of signaling proteins and reduces the levels of CtrA~P. Steady-state distributions of CtrA~P concentration along the long axis of the cell is shown in wildtype conditions (i) and with the addition of a new protein (purple) that competes for PopZ binding sites in the new pole microdomain (ii). The pie charts (right) show the relative occupancy of PopZ binding sites by CckA, ChpT, CtrA, and the competitive binder (blue, orange, green, purple). With the addition of the competitive binder the number of ChpT and CtrA molecules within the microdomain is reduced ((ii), pie chart). Under this condition, the total amount of CtrA~P molecules is reduced by 42% and the gradient shape becomes less steep. (F) The predivisional cell maintains a sublinear CtrA~P gradient. Starting with 10,000 unphosphorylated CtrA molecules, within the course of 8 minutes of simulation, 63% of CtrA molecules are phosphorylated and a gradient of CtrA~P is established. CtrA~P maintains a similar sublinear distribution over time, as shown at 10 equally spaced time intervals (light green to blue lines). In each interval at least 70% of all CtrA~P molecules are found in newly formed daughter swarmer cell (at most 2 gm away from the new pole).

To determine what aspects of sequestration within the PopZ microdomain were most important in controlling the distribution of CtrA~P, we performed a global sensitivity analysis of our model parameters (SI). In particular, we investigated how the amplitude (total CtrA~P) and shape (CtrA~P distribution between the nascent daughter cells) of the phospho-gradient were affected by changes in the binding affinity to each of the new and old pole microdomains, and the rates of phosphotransfer and the diffusion coefficients inside and outside the microdomains (Figure S6A). The amplitude of CtrA~P was highly sensitive to alterations in the K_D_ of binding between CckA and the new pole microdomain as well as the phosphotransfer rate from CckA to ChpT (Figure S6A upper heatmap). Changing the K_D_ between CckA and the new pole microdomain alters the concentration of CckA, and because CckA’s kinase activity increases nonlinearly with increasing concentration (Mann et al., 2016), this greatly increases the rate at which phosphate is introduced into the pathway (Figure 5B). Meanwhile, faster phosphotransfer rates between CckA and ChpT and between ChpT and CtrA increase the fraction of CtrA~P molecules that is generated at the new pole microdomain, near the CckA source of phosphate. The distribution of CtrA~P was highly sensitive to changes in ChpT and CtrA binding to the microdomain at the new pole, as well as to CtrA diffusion in the body of the cell (Figure S6A lower heatmap). Weakening the K_D_ of ChpT or CtrA for the new pole microdomain led to a shallower gradient across the cell, with a larger contribution from ChpT. Strong interactions between both proteins and the new pole were critical to achieve a CtrA~P distribution that, *in-silico*, recapitulates the transcriptional measurements (Figure 5C). These results suggest that the spatial position of the phosphotransfer between ChpT and CtrA can modulate the shape of the gradient. Additionally, it was critical that CtrA diffusivity in the body not be substantially higher than the effective kinetics of phosphotransfer, as this would disrupt any gradient. To illustrate the molecular transfer mechanism underlying these changes in steady-state output, we modeled the rate and location of phospotransfer events within the CtrA~P pathway. For the “wild-type” system, we observed that more than 95% of forward transfer events (from CckA~P to ChpT and from ChpT~P to CtrA) occur within the new pole microdomain, while back transfer events (from CtrA~P to ChpT and from ChpT~P to CckA) occur everywhere away from the new pole (Figures 5D and S6B), highlighting the importance of coordinated transfer in this region (Figure S6A). By contrast, for an *in silico* mutant with weakened binding affinity between ChpT and the microdomain (KD increased from 10 nM to 10 μM), phosphotransfer coordination was lost and 60% of ChpT~P to CtrA transfer events occurred outside of the new pole hub, giving a molecular mechanism for the less pronounced gradient shown in Figure 5C.

Our single-molecule measurements showing protein exclusion from the pole indicate that that client permeability of the microdomain can be modulated by PopZ and client copy number. One potential contribution to permeability may be client density due to competition for free volume or binding sites within the PopZ matrix. We chose such occupancy-based regulation to investigate how universal regulation of microdomain permeability would alter the output of the CtrA pathway. To model this competition mechanism, we introduced a separate cytosolic protein with high binding affinity for the microdomain while keeping the binding affinities of CckA, ChpT and CtrA to the microdomain constant (SI). The simulated reduction in the microdomain accessibility led to a 50% decrease in the relative proportion of ChpT and CtrA inside the poles, reducing the efficiency of the signaling pathway and overall CtrA~P levels while maintaining a roughly similar gradient shape within the cell body (Figure 5E). Thus, our simulations show that both specific binding interactions and overall PopZ microdomain permeability play key roles in the formation of a CtrA~P gradient in terms of both amplitude and shape.

CtrA molecules are cleared from the cell during differentiation and resynthesized in predivisional cells over the course of 60 minutes (Domian et al., 1997; Quon et al., 1996). We used our model to determine how the CtrA~P gradient evolves from newly synthesized CtrA molecules in predivisional cells. We found that for a given number of total CtrA molecules, CtrA is rapidly phosphorylated and that the CtrA~P distribution amplitude reaches steady state within five minutes of simulation time (Figure 5F). The nonlinear gradient in CtrA~P distribution is established immediately at the beginning of the simulation and is maintained until reaching its steady state. Thus, the quasi-steady-state CtrA~P gradient is established early in predivisional cells as CtrA molecules are synthesized, robustly priming the nascent daughter cells for asymmetric cell division.

## DISCUSSION

Membrane-enclosed compartments are an effective means for concentrating functional complexes to isolated environments within the cell across all kingdoms of life. Alternatively, membrane-less compartments have emerged as a widespread organizational unit. These include membrane-less organelles in eukaryotes (Alberti, 2017; Banani et al., 2017; Shin and Brangwynne, 2017) and protein-encapsulated microcompartments in prokaryotes and archaea (Kerfeld and Erbilgin, 2015). Eukaryotic membrane-less compartments are held together by weak interactions between their components that form agglomerated condensates (Molliex et al., 2015; Nott et al., 2015). Unlike structurally defined macromolecular complexes the size and internal organization of such condensates are both heterogeneous and dynamic (Shin and Brangwynne, 2017). Here, we showed that the *Caulobacter* PopZ microdomain, which lacks a membrane enclosure and does not appear to be encapsulated by a protein shell, differs from bacterial microcompartments while sharing key qualities with eukaryotic condensed compartments. Most significantly for function, the PopZ microdomain size is dynamic, client entry into the microdomain is selective, and clients dynamically exchange with the cytosol. We further showed that these properties facilitate accelerated phosphotransfer reactions and modulation of the amplitude and distribution of CtrA~P in the predivisional cell.

### Organization and selective permeability of the space-filling PopZ microdomain

PopZ self-assembles to organize a space-filling microdomain (Bowman et al., 2013; Gahlmann et al., 2013; Ptacin et al., 2014) that coordinates a suite of cell polarity functions including the phosphotransfer cascade controlling CtrA activity. Whereas PopZ itself is cytoplasmic, both transmembrane and cytosolic client proteins are (to our imaging resolution) uniformly distributed adjacent to and throughout the space-filling PopZ microdomain for transmembrane and cytosolic proteins, respectively (Figures 3 and 4) (Ptacin et al., 2014). PopZ is recruited to the membrane by direct binding to several membrane-bound factors, and mislocalization of these factors can seed ectopic PopZ microdomains (Holmes et al., 2016; Perez et al., 2017). Thus, the shape of the PopZ microdomain appears to be defined by a condensation and wetting process that balances cohesive forces (self-interaction) and adhesive forces (membrane-protein interactions), similar to that observed for tau protein droplets on microtubules (Hernandez-Vega et al., 2017). This balance allows PopZ to lie across a substantial membrane surface area while still creating a large but porous space-filling volume, bridging the membrane and the cytoplasm and jointly concentrating both membrane and cytoplasmic proteins.

The PopZ microdomain selectively captures and concentrates target proteins at the cell pole, including the three CtrA pathway proteins. Proteins are recruited to the PopZ microdomain either by direct binding to PopZ, as shown for at least nine client proteins (Holmes et al., 2016), or indirectly by binding to one or more proteins which bind PopZ, as shown for the membrane protein DivJ (Perez et al., 2017) and the cytosolic protein CtrA (Figures 1D, 4C-E, and 6). Whereas large molecules and complexes such as DNA and ribosomes (Bowman et al., 2008; Ebersbach et al., 2008) are excluded from the microdomain, we have shown that small proteins lacking a microdomain binding partner, such as free eYFP and fPIF-eYFP, cannot penetrate the PopZ microdomain despite being similar in size and charge to proteins that do enter (Figures 4H and 6, Table S1). Notably, when the PopZ microdomain is drastically expanded by exclusively overexpressing *popZ*, fluorescent proteins, including eYFP, can enter the microdomain (Ebersbach et al., 2008; Laloux and Jacobs-Wagner, 2013), suggesting that changes in the microdomain such as occupancy can regulate its permeability prosperities. Indeed, studies of eukaryotic condensates have identified multiple factors that affect permeability, including occupancy by additional binding partners (Banani et al., 2016), charged proteins (Nott et al., 2016), or RNAs (Wei et al., 2017), as well as environmental properties including viscosity, dielectric constant, and scaffold-scaffold interaction strength within the condensate. Our observation that eYFP and fPIF-eYFP cannot enter the wildtype microdomain interior even transiently (Figure 4H) demonstrate that even without membrane barriers or a protein shell, the PopZ microdomain forms a compartmentalized volume of strictly defined composition. Our results indicate that the membrane-PopZ and the PopZ-cytoplasm interfaces are critical junction points (for phosphate transfer from CckA, Figure 4B, and escape of CtrA, Figure 4D). Therefore, the protein composition and structure at these interfaces can be expected to be particularly important for microdomain function, and in future work, it will be critical to determine whether the already distinct “microenvironment” created by PopZ contains further distinct “nanoenvironments” in these regions.

### Efficient phosphotransfer within the PopZ microdomain

Joint sequestration of CckA, ChpT, and CtrA acts to locally enhance pathway activity within the PopZ microdomain (Figures 4-6). We previously reported *in vitro* reconstitution experiments showing that high densities of CckA on liposomes enhance its autokinase activity (Mann et al., 2016). In keeping with these results, CckA functions as a kinase *in vivo* solely when sequestered at the new pole of predivisional cells (Iniesta et al., 2010; Jacobs et al., 2003; Jacobs et al., 1999), where the new pole-specific factor DivL is recruited to allosterically modulate CckA kinase activity (Childers et al., 2014; Iniesta et al., 2010; Tsokos et al., 2011). CckA positioned elsewhere in the cell primarily acts as a phosphatase (Iniesta et al., 2010; Mann et al., 2016). Indeed, the dependence of CckA autokinase activity on concentration implies that the ~500-1,000x higher CckA density achieved within the new pole is necessary for CckA to act as a phosphate source (SI) (Mann et al., 2016). Further, the elevated concentrations of all three proteins increase the probability of intermolecular binding and phosphate transfer events between CckA and ChpT, and ChpT and CtrA via mass action (Figure 6C), and the vast majority of phosphotransfer events take place within the microdomain (Figure 6D). Similarly, a model for the function of the chromosome partitioning protein ParA (Ptacin et al., 2014) suggests that high ParA concentration and thus an increased rate of homodimerization within the PopZ microdomain is critical for chromosome segregation. More broadly, membrane-less compartments are emerging as an important mechanism for creating specialized condensed signaling environments (Chong and Forman-Kay, 2016; Li et al., 2012; Su et al., 2016). This function is reminiscent of scaffolding molecules that bind multiple signaling proteins simultaneously to regulate selectivity and adaptation to stress (Atay and Skotheim, 2017; Good et al., 2011). Quantitative characterization of the unique properties of each architecture would provide insight into the advantages of using either subcellular organization strategy.

**Figure 6.**
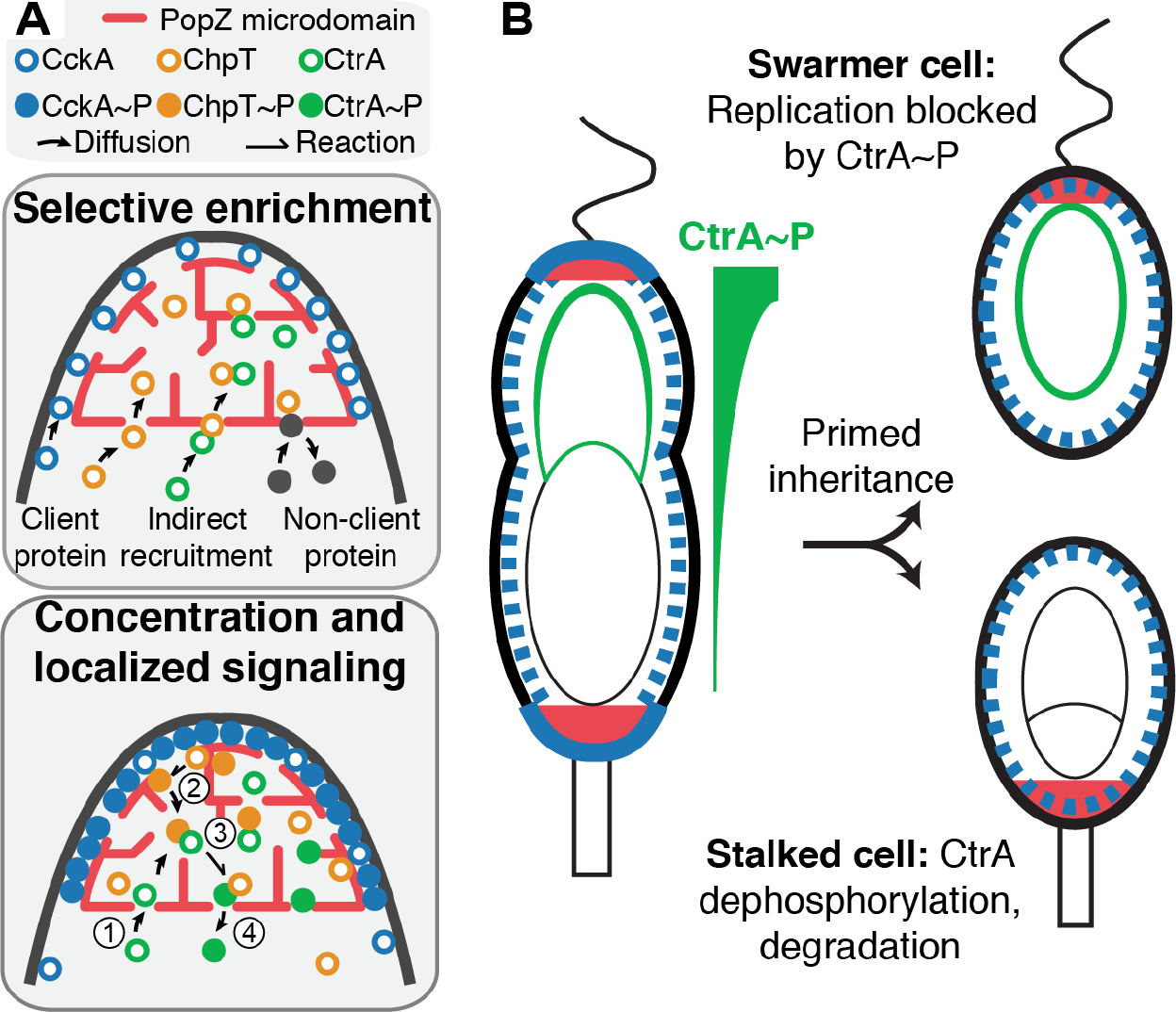
A transcriptional gradient propagates from the PopZ microdomain at the new pole, in which cell fate signaling flow is altered. **(A)** Specialized properties of the new pole PopZ microdomain establish a local zone of differential signaling flow. Selective entry of client proteins is permitted via direct interaction with PopZ or by interaction with a PopZ client (indirect recruitment), whereas non-client proteins cannot enter. Proteins localized to PopZ are concentrated in the microdomain due to extended dwell times relative to the cell body. Increased concentration of CckA-ChpT-CtrA pathway components leads to a local increase in CckA kinase activity and thus CtrA~P output. The numbers represent stages of the phosphotransfer pathway; 1) recruitment of proteins to the microdomain, 2) complex formation and transfer of phosphate from CckA~P to ChpT, 3) complex formation and transfer of phosphate from ChpT~P to CtrA, and 4) exit of CtrA~P from the microdomain, establishing a local source of CtrA~P. **(B)** CckA acts as a kinase while accumulated at the new cell pole (blue bar) while its phosphatase activity dominates when it is diffusely localized (blue dashes) in the rest of the cell. This localized source of kinase signaling flow, in conjunction with slowed CtrA~P diffusion across the long axis of the cell, ultimately leads to a gradient of CtrA~P that primes each daughter cell for a distinct cell fate. High CtrA~P blocks DNA replication in the swarmer progeny, whereas CckA phosphatase activity in the stalked progeny leads to dephosphorylation and ultimately degradation of CtrA, permitting DNA replication and differential readout of the chromosome in the two progeny cells. The theta structure represents the partially replicated chromosome in the predivisional cell and stalked progeny, while the swarmer cell has a single circular chromosome. Green shading along the chromosome represents binding by CtrA~P to the origin, which forms a gradient in predivisional cells, is uniform in swarmer progeny, and is absent in stalked progeny.

Our results show that in addition to concentrating proteins, the intrinsic character of the PopZ microdomain slows turnover between the pole and the cell body. The observed rates of ChpT and CtrA exit from the pole, on the order of 100 ms (Figure 4), are several times slower than would be expected without sequestration within the microdomain. This slow turnover results from a combination of binding to PopZ and other polar proteins (as implied by localization to the pole), and to percolation through the microdomain volume (reflected by the slowed polar diffusion of CtrA and to a lesser extent, ChpT, Figure 4). Similarly, while CckA still diffuses freely (though more slowly) at the polar membrane, binding interactions slow the exchange of CckA between the polar microdomain and the rest of the cell (Figure 3D). Tracking experiments with gold nanobead labels in mammalian cells have shown that physical obstruction by cortical actin networks causes transmembrane proteins to exhibit “hop diffusion” between nanoscale domains on the sub-ms timescale (Fujiwara et al., 2016). While such phenomena are below the resolution of our experiments, obstruction by PopZ polymers close to the membrane may be partly responsible for slowed CckA motion at the poles and could play a role in the organization of membrane proteins within the microdomain.

### A sequestered phospho-signaling cascade drives a gradient of active CtrA

Sensitivity analysis in our reaction-diffusion model showed that sequestration of all three proteins within the new pole microdomain is critical for establishing a CtrA~P gradient (Figure S6A). Both ChpT and CtrA must be localized at high density for rapid phosphotransfer within the pole, while allowing ChpT to transfer its phosphate to CtrA outside of the microdomain would reduce or eliminate the gradient (Figures 6C-E). Critically, we found replacing *ctrA* with its phosphomimetic mutant *ctrA*(*D51E*), which is active independent of the polar signaling cascade, abolished the gradient of transcriptional activity and disrupts normal cell cycle progression (Figure 2B).

Spatial separation of opposing enzymes in activation–deactivation cycles of protein modification is necessary for maintaining a protein activity gradient (Brown and Kholodenko, 1999). Because the phosphotransfer cascade that activates CtrA is conducted via the intermediate cytosolic protein ChpT, the requirements for maintaining a steady state gradient of CtrA~P are more complex than a canonical gradient established by two factors. In systems with one phosphotransfer step, in which the phosphatase is far from saturating, the gradient decays almost exponentially as a function of the ratio between diffusion of signal and phosphatase rate (Barkai and Shilo, 2009; Tropini et al., 2012; Wartlick et al., 2009), while in systems with a cascade of phosphotransfer steps, co-localization of the pathway components plays a key role in gradient shape (Kholodenko et al., 2010), such as created by scaffolding proteins (Good et al., 2011). Here we demonstrated that a bacterial microdomain exhibits a similar function to eukaryotic scaffolds by sequestering all members of the CtrA activation pathway and effectively combining the forward phosphotransfer activities of CckA and ChpT into one localized enhanced source, generating a gradient of CtrA~P across the entire cell (Figures 5 and 6).

### A stable CtrA~P gradient facilitates asymmetric division

*Caulobacter* asymmetric cell division yields daughter cells that exhibits different genetic readouts despite having identical genomes (McAdams and Shapiro, 2009). Differences in gene expression profiles stem from the asymmetric inheritance of the CtrA~P transcription factor controlling the expression of over 100 cycle-regulated genes, many of which encode swarmer cell-specific functions (Laub et al., 2002; Laub et al., 2000). We showed that the CtrA~P distribution is not only skewed toward the new pole but decreases rapidly away from the new pole forming a stable sublinear CtrA~P gradient (Figures 2 and 5A). In our simulations, the CtrA~P gradient is formed rapidly and maintains its shape, showing that this output is reached quickly and reliably (Figure 5F). We demonstrated using RT-qPCR that timing of the CtrA-regulated gene expression can be modulated by altering the position of the gene on the chromosome (Figure 2A). These results suggest that the induced asymmetry in CtrA~P concentration before division may be used by the cell as regulator of gene expression timing. Indeed, the relative positions of CtrA regulated genes is conserved across alpha-proteobacteria (Ash et al., 2014) and movement of the *ctrA* transcriptional unit from the vicinity of the origin to the vicinity of the termini disrupts normal cell cycle progression (Reisenauer and Shapiro, 2002).

The skewed inheritance of CtrA~P, primed by the gradient, is critical for the differential ability of the two daughter cells to initiate chromosome replication. The daughter stalked cell, in which inherited CtrA~P levels are low, initiates a new round of DNA replication immediately after cytokinesis, whereas the daughter swarmer cell, with high levels of CtrA~P, is arrested in G1 phase until it differentiates into a stalked cell (Quon et al., 1996; Quon et al., 1998). In its phosphorylated form, CtrA~P inhibits initiation of DNA replication by binding to specific sites in the origin region (Quon et al., 1998). Indeed, it was previously suggested that CtrA~P is skewed towards the new cell pole based on the observation that additional replication events happened more often at the stalked cell pole when normal cytokinesis and cell cycle progression was disrupted (Chen et al., 2011). Maintaining a sharp gradient of CtrA~P concentration in the predivisional cell therefore facilitates the robustness of asymmetric division by controlling both differential gene expression and replication initiation.

### Outlook

It was recently shown that signaling proteins in eukaryotic cells can not only spontaneously separate into liquid-like agglomerated condensates with distinct physical and biochemical properties, but that these condensates can selectively exclude proteins with antagonist function (Su et al., 2016). Our work highlights how selective sequestration of a signaling pathway within a bacterial membrane-less microdomain provides an environment for accelerated biochemistry that directs signaling across the entire cell with exquisite spatial control culminating in differential activity of a key transcription factor that controls an asymmetric division. Although we do not know the physical nature of the microdomain, nor whether the microdomain separates into a liquid- or gel-like phase, its major constituent, PopZ, is a low complexity protein that forms a mesh-like structure *in vitro*, and self assembles into a distinct cytoplasmic microdomain *in vivo*. The selective permeability of the PopZ microdomain is reminiscent of the eukaryotic nuclear pore complex (NPC), which mediates active transport between nucleus and cytoplasm (Knockenhauer and Schwartz, 2016). The NPC gains a remarkable sorting selectivity from disordered domains that come together in a meshwork and selectively bind to pockets in the proteins that are marked for transport, suggesting a possible functional parallel to the organization and function of the PopZ microdomain (Fu et al., 2017). Notably, similarities have been drawn between the NPC low complexity domains and P granules, a well-studied membrane-less organelle (Schmidt and Gorlich, 2016). Thus, further structural and functional characterizations of the bacterial PopZ microdomain will be pivotal for revealing fundamental principles for subcellular organization on the mesoscale.

## ACKNOWLEDGMENTS

We thank Justin W. Kern for help with plasmid design and construction, Michael D. Melfi for providing a RT-qPCR protocol, Alberto Lovell and Michael R. Eckart at Stanford PAN Facility for their support in conducting RT-qPCR, Jared M. Schrader for sharing unpublished data on mRNA half-life in *Caulobacter*, Andrew Olson at the Stanford Neuroscience Microscopy Service for assistance with photobleaching experiments, as well as Allison Squires for critical feedback on the manuscript. We also thank Harley H. McAdams for helpful discussions on modeling of signal transduction, Darshankumar Pathak and all members of the Shapiro and Moerner labs for helpful discussions throughout the project. We acknowledge support from the Gordon and Betty Moore Function GBMF 2550.03 to Life Sciences Research Foundation [to K.L.], the Weizmann Institute of Science National Postdoctoral Award Program for Advancing Women in Science [to K.L.], and from NIGMS of the National Institutes of Health under award numbers T32GM007276 [to T.H.M], R01-GM086196 [to W.E.M. and L.S.], R35-GM118067 [to W.E.M.], and R35-GM118071 [to L.S.]. L.S. is a Chan Zuckerberg Biohub Investigator.

The content is solely the responsibility of the authors and does not necessarily represent the official views of the National Institutes of Health.

